# Pre-metazoan origin of animal miRNAs

**DOI:** 10.1101/076190

**Authors:** Jon Bråte, Ralf S. Neumann, Bastian Fromm, Arthur A. B. Haraldsen, Paul E. Grini, Kamran Shalchian-Tabrizi

**Affiliations:** Centre for Epigenetics, Development and Evolution (CEDE) and Centre for Integrative Microbial Evolution (CIME), Section for Genetics and Evolutionary Biology (EVOGENE), University of Oslo, Norway.; Institute of Clinical Medicine, Department of Immunology, University of Oslo, Norway.; Department of Tumor Biology, Institute for Cancer Research, Norwegian Radium Hospital, Oslo University Hospital, Norway.

**Keywords:** miRNA, animal evolution, *Sphaeroforma*, Ichthyosporea, Holozoa

## Abstract

microRNAs (miRNAs) are integrated parts of the developmental toolkit in animals. The evolutionary history and origins of animal miRNAs is however unclear, and it is not known when they evolved and on how many occasions. We have therefore investigated the presence of miRNAs and the necessary miRNA biogenesis machinery in a large group of unicellular relatives of animals, Ichthyosporea. By small RNA sequencing we find evidence for at least four genes in the genus *Sphaeroforma* that satisfy the criteria for the annotation of animal miRNA genes. Three of these miRNAs are conserved and expressed across sphaeroformid species. Furthermore, we identify homologues of the animal miRNA biogenesis genes across a wide range of ichthyosporeans, including *Drosha* and *Pasha* which make up the animal specific Microprocessor complex. Taken together we report the first evidence for *bona fide* miRNA genes and the presence of the miRNA-processing pathway in unicellular Choanozoa, implying that the origin of animal miRNAs and the Microprocessor complex predates multicellular animals.

## Introduction

microRNAs (miRNAs) are short (20-26 nucleotides) RNA molecules that are involved in gene regulation (Fromm, 2016). They are found in a wide variety of organisms including plants, viruses and animals but current evidence does not support a common evolutionary history (Tarver *et al.*, 2012; Cerutti and Casas-Mollano, 2006; Shabalina and Koonin, 2016). In animals, miRNAs are essential components of the gene regulatory machinery and highly important for processes like cell differentiation and development Mello and Conte, 2004. Furthermore, because of the increase of distinct miRNA families during animal evolution (combined with lineage specific expansions and rare losses), miRNAs have been linked to the evolution of morphological complexity in animals (Berezikov, 2011; Wheeler *et al.*, 2009; Kosik, 2010; Fromm *et al.*, 2013; Erwin *et al.*, 2011; Philippe *et al.*, 2011). Mainly based on the absence of miRNAs in single-celled ancestors, it has even been proposed that miRNAs were one of the key factors that allowed multicellular animals to evolve (Grimson *et al.*, 2008; Wheeler *et al.*, 2009; Mattick, 2004).

Generally, the primary transcripts of miRNA genes (pri-miRNA) can be very long RNA molecules that contain hairpin structures (Axtell et al. 2011; Ameres & Zamore 2013; Chang et al. 2015). In animals, the so-called Microprocessor complex (Han *et al.*, 2006), comprising the RNase III enzyme Drosha and the double-stranded RNA (dsRNA) -binding protein Pasha (known as DGCR8 in humans), subsequently excises the hairpin structures (creating the pre-miRNA). In contrast, plants do not possess Drosha or Pasha proteins; here, several copies of the nuclear RNase III protein Dicer fulfills the role of the Microprocessor. In both animals and plants, the loop region of the hairpin pre-miRNA is subsequently removed by Dicer proteins, albeit dicing takes place in the cytoplasm and nucleus, respectively, resulting in a 20-26 nt dsRNA molecule. Typically, only one of these strands (known as the mature strand) is then loaded into the RNA-induced silencing complex (RISC), while the other strand (the passenger, or star strand) is degraded. Gene regulation takes place via the RISC and the central component, the Argonaute (AGO) protein that binds the mature miRNA. Target messenger RNAs (mRNAs) are identified by basepairing between the loaded miRNAs and the mRNA, and translation of the mRNA is either fully blocked or down regulated.

In animals, miRNA genes have been found even in the early diverging animal lineages such as sponges (Grimson *et al.*, 2008; Robinson *et al.*, 2013; Liew *et al.*, 2016; Wheeler *et al.*, 2009). In fact, only two animal lineages are known to be missing miRNAs, the ctenophores and Placozoa (Maxwell et al. 2012; Grimson et al. 2008; Moroz et al. 2014). However, due to the lack of homologous miRNA sequences in the basal branching animals, it has been hypothesized that miRNA genes in animals arose several times independently (Tarver *et al.*, 2012). Although, except from one study sequencing the small RNA content of the choanoflagellate *Monosiga brevicollis* (Grimson et al. 2008), almost nothing is known about the presence of miRNA genes, or the biogenesis machinery, in the unicellular relatives of animals.

Here, we therefore investigate whether miRNA genes coevolved with animal multicellularity or if these genes were present already in the unicellular ancestors of animals, the choanozoans. We address this question by deep sequencing of smallRNAs from species belonging to Ichthyosporea; a basal group at the split between animals and fungi, and search for the animal miRNA biogenesis machinery in all unicellular relatives of animals. Our results show the existence of six miRNA genes in *Sphaeroforma* that fulfill all criteria for the annotation of metazoan miRNAs (Fromm *et al.*, 2015). Three of the miRNAs are highly conserved and expressed across three species, which strongly indicate that the genes have important and similar function. In addition, we show that the genomes of several species of Ichthyosporea, covering the genera *Sphaeroforma*, *Creolimax*, *Pirum*, *Abeoforma* and *Amoebidium* contain homologs of the animal *Pasha* and *Drosha* genes as well as *Dicer*, *Exportin 5* and *AGO*.

Altogether we present an evolutionary scenario where the common ancestor of Choanozoa and animals (Holozoan ancestor) already possessed the animal miRNA biogenesis pathway, including *Drosha* and *Pasha*, as well as *bona fide* miRNAs. This implies that both the animal miRNAs and the Microprocessor pre-date the origin of animals and that the last common ancestor of animals likely used miRNAs for gene regulation.

## Results

### Sphaeroforma species have bona fide miRNA genes

Initially, our small RNA sequencing and bioinformatics pipeline identified six miRNA genes in *Sphaeroforma arctica*(Figure 1 and Supplementary File 1). Five of the miRNAs fulfilled all of the recently proposed criteria for the annotation of miRNA genes (Fromm *et al.*, 2015); at least two 20-26 nt RNA strands are expressed from each of the two arms that are derived from a hairpin precursor with 2-nt offsets (note that the 1nt Drosha-offset of Sar-Mir-Nov-3 and the 1nt Dicer-offset of Sar-Mir-Nov-5 are both likely due to the low number of reads in general and correspondingly lower read-counts of the star sequence). Further, the reads on both arms show no or very little variation of their 5’ start positions (5’ homogeneity) and we observe at least 16 complementary nucleotides between the two arms. We include here Sar-Mir-Nov-6, although the star strand is not expressed and thus Dicer or Drosha offsets cannot be assessed because it shows high expression, a good structure and is located close to Sar-Mir-Nov-5 forming a polycistronic miRNA. The loop sizes are quite variable and Sar-Mir-Nov-5 has the smallest loop size of 36 nts. Sar-Mir-Nov-1, 2 and 4 are slightly larger with 42, 41 and 43 nts respectively, while Sar-Mir-Nov-3 and Sar-Mir-Nov-6 are considerably bigger with loop sizes of 59 and 60 nucleotides.

**Fig 1.**
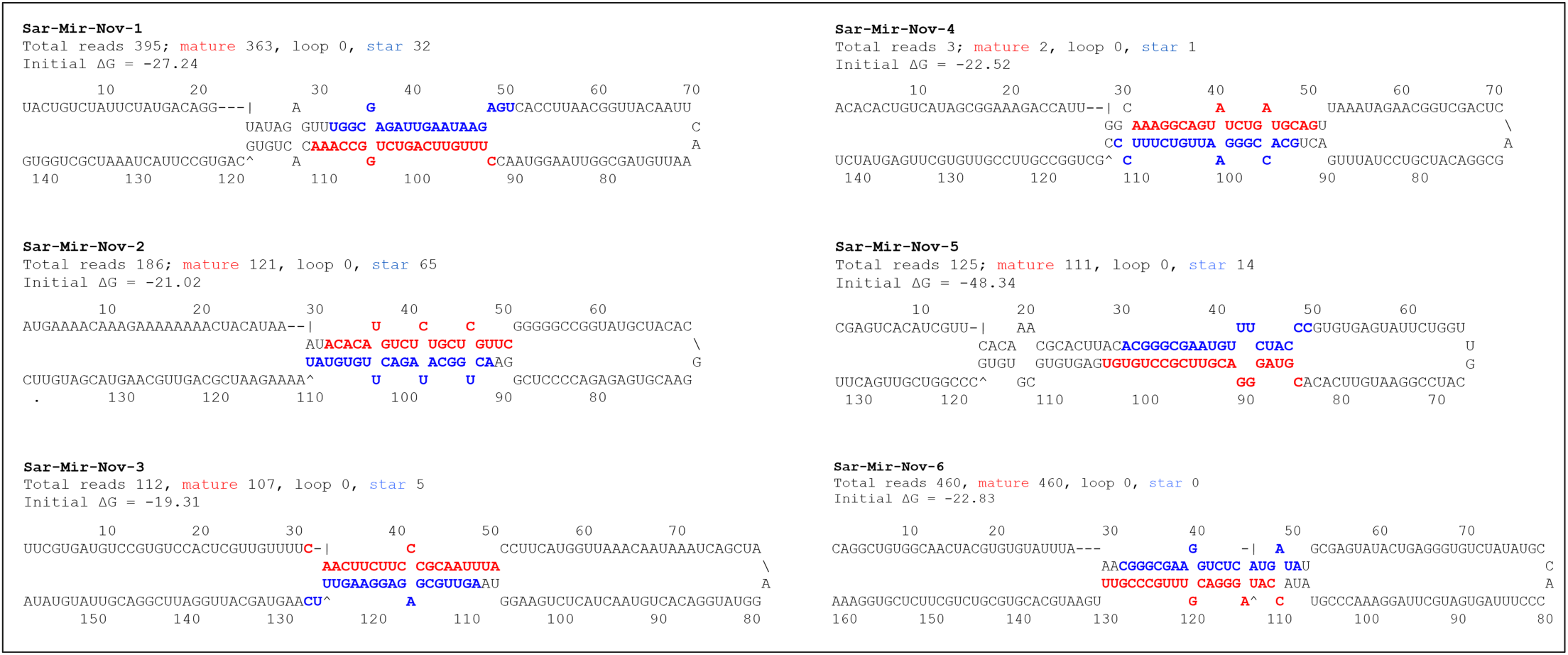
Novel Ichthyosporean miRNAs. The six miRNA structures as they are identified in *S. arctica*. Red font indicates the mature strand and blue font indicates the star strand. The secondary structures shown here are constrained to not show base pairing in the loop regions.

The expression of the primary miRNA transcripts is confirmed by transcription start site (TSS) and mRNA sequencing (Figure S1). Using the same pipeline on *S. sirkka* we identified four of the *S. arctica* miRNAs (Table 1 and Supplementary File 1). No other additional miRNAs were found in *S. sirkka*. Because there is no genome available for *S. napiecek* we could not run the miRNA prediction tools as for the other species. But we could identify five of the *S. arctica* pre-miRNAs among the expressed mRNAs (even though the pri-miRNA is transcribed by RNA polymerase II, and hence is poly-adenylated, it might not be present among the mRNAs because of rapid processing by Drosha and Dicer). These pre-miRNAs also had small RNA reads mapping to the same locations as the mature and star strands in *S. arctica*, except for the star region of Sar-Mir-Nov-3 and 5. The pre-miRNAs were highly conserved between the three species with most mutations in the loop region which does not affect the gene targeting. But there were also a few minor differences in the mature and star strands, most notably in Sar-Mir-Nov-2. None of the novel miRNAs had any detectable homologs outside of Ichthyosporea.

**Table 1.**
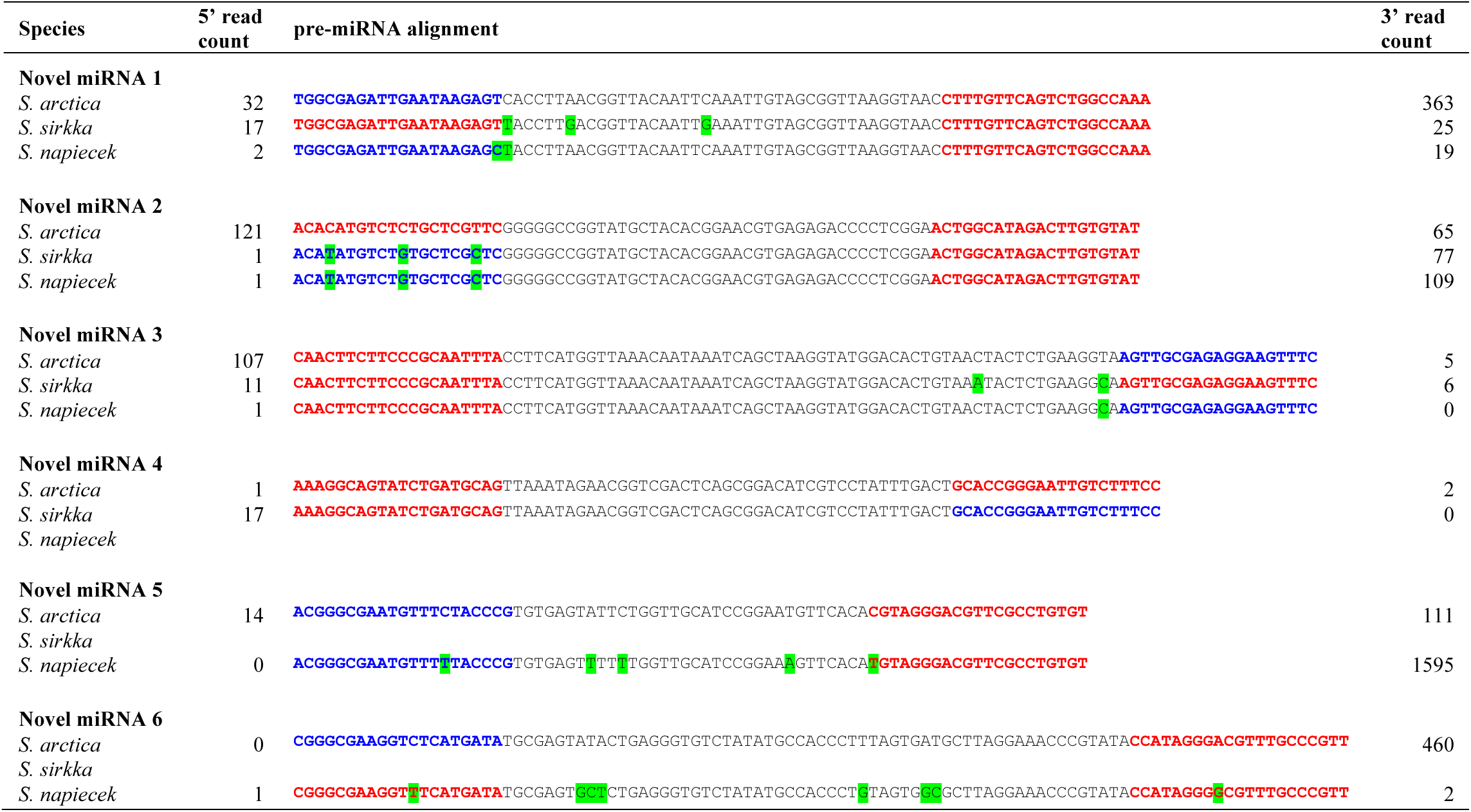
Conservation of miRNAs across *Sphaeroforma*. Alignment of the pre-miRNA regions shows high degree of conservation across *S. arctica*, *S. sirkka* and *S. napiecek*. Mutations in the pre-miRNAs compared with *S. arctica* are highlighted in green. The mature and star strands are highlighted with red and blue font respectively. In cases where the 5p:3p mapping ratio is below 2, co-mature strands are indicated as defined by Fromm *et al*. (2015).

The miRNA genes are either located in intergenic regions or in the introns and UTRs of protein-coding genes (Figure S1). In *S. arctica* Sar-Mir-Nov-1, 3, 5 and 6 are encoded in intergenic regions and transcribed as > 1kb unspliced pri-miRNA fragments with a well-defined TSS, except for Sar-Mir-Nov-3 which is lacking a TSS but this is probably an artifact of the genome assembly. Interestingly, the region covering the Sar-Mir-Nov-5 pri-miRNA is transcribed also from the opposite strand which could represent a potential novel long non-coding RNA with binding capacity to the miRNA (an miRNA sponge) as no protein-coding gene is annotated in the region. Sar-Mir-Nov-2 and Sar-Mir-Nov-4 are encoded within protein coding genes, in the intron and the 3’-UTR respectively (Figure S1). These miRNAs are not transcribed from a separate TSS, but instead co-transcribed with the protein-coding gene and subsequently processed into the mature miRNAs. It is worth noting that these miRNAs are located within the *Drosha* and *Argonaute* genes, indicating a possible co-evolution of the miRNAs and the miRNA processing genes.

### The S. arctica genome contains both plant- and animal-type miRNA targets

In animals, mRNA targets in 3’-UTRs, so called miRNA response elements (MREs) are usually recognized by short seed regions (7 nts) of the mature strand (Bartel, 2009). As a consequence, target prediction is highly uncertain. Showing that putative miRNA target sequences are conserved in homologous target genes between species increases the probability that a true miRNA target is found, but needs to be verified experimentally. However, such conserved target sequences are not available from *Sphaeroforma* and thus could not be included as input to the target prediction algorithm. Still, many potential target genes in the *S. arctica* genome were identified for all the miRNAs using the TargetScan program (Agarwal *et al.*, 2015) with a context+ score cutoff of −0.2 as in Jacobsen *et al.* (2013).

We additionally searched for plant-like miRNA targets in the *S. arctica* transcriptome using the TAPIR web server (Table S2; Bonnet *et al.*, 2010). Plant-like targets require a near-perfect basepairing along the whole length of the mature miRNA sequence and are thus much less likely to be an artifact. For Sar-Mir-Nov-3 we identified one gene with a plant-like target mode which is clearly homologous to an ABC transporter. Sar-Mir-Nov-4 also targets a single gene. This gene has no conserved domains but a weak Blast hit against the RED1 gene from yeast, a gene required for chromosome segregation during meiosis Thompson and Roeder, 1989. For Sar-Mir-Nov-5 and 6 we identified three target genes with a plant-like target mode. These three genes are likely paralogs as they share significant sequence similarity (including 13 additional homologs in the *S. arctica* genome, data not shown). These three genes are all zinc-finger transcription factors.

### Ichthyosporea species harbor key elements of the animal miRNA processing machinery

We used reciprocal Blast searches and domain detection tools to identify genes related to the Microprocessor and RNAi machinery. We used all available Choanozoa genomes and transcriptomes, including the new transcriptomes generated in this study. The genes we searched for were *Argonaute*(*AGO*), *Dicer*, *Drosha*, *Pasha*, and *Exportin 5*(*XPO5*). Besides *XPO5*, which was detected in all opisthokont lineages, the choanoflagellates and filastereans seems to be lacking all key genes involved in miRNA processing (Table 2). No Argonaute of the *AGO* subfamily (with the signature PAZ and PIWI domains) was detected, likewise no *Dicer* or *Drosha* gene, nor *Pasha* was detected. Importantly however, these genes were all detected in most of the ichthyosporean species (Table 2 and S3). The annotated *Dicer*/*Drosha* genes contained two consecutive RNaseIII domains, which is a unique feature of *Dicer* and *Drosha* genes (Supplementary File 2 and Supplementary Information) revealing that both types of genes may be present (a few short genes identified as either *Dicer* or *Drosha* by the Blast annotation pipeline covered only a single RNase III domain likely due to incomplete assembly).

**Table 2.** Identification of genes related to the miRNA biogenesis machinery in animals based on reciprocal Blast analyses, domain annotation and phylogenetic analyses of gene families. Shaded rows indicate results new in this study.

|  | Argonaute | Exportin 5 | Dicer | Drosha | Pasha |
| --- | --- | --- | --- | --- | --- |
| **Porifera** |  |  |  |  |  |
| *Amphimedon queenslandica* | + | + | + | + | + |
| *Acropora digitifera* | + | + | + | + | + |
| **Choanoflagellata** |  |  |  |  |  |
| *Acanthoecca spectabilis* | - | - | - | - | - |
| *Acanthoecca sp.* | - | - | - | - | - |
| *Monosiga brevicollis* | - | + | - | - | - |
| *Salpingoeca pyxidium* | - | + | - | - | - |
| *Salpingoeca rosetta* | - | + | - | - | - |
| **Filasterea** |  |  |  |  |  |
| *Capsaspora owczarzaki* | - | + | - | - | - |
| *Ministeria vibrans* | - | - | - | - | - |
| **Ichthyosporea** |  |  |  |  |  |
| *Abeoforma whisleri* | + | + | + | + | + |
| *Amoebidium parasiticum* | + | + | + | + | + |
| *Corallochytrium limacisporum* | - | + | - | - | - |
| *Creolimax fragrantissima* | + | + | - | + | + |
| *Ichthyophonus hoferi* | + | + | + | - | - |
| *Pirum gemmata* | + | - | + | + | + |
| *Sphaeroforma arctica* | + | + | + | + | + |
| *Sphaeroforma napiecek* | + | + | + | + | + |
| *Sphaeroforma sirkka* | + | + | + | + | + |
| *Sphaerothecum destruens* | - | + | - | - | - |
| **Amoebozoa** |  |  |  |  |  |
| *Dictyostelium discoideum* | + | + | + | - | - |
| **Nucleariidae** |  |  |  |  |  |
| *Fonticula alba* | - | + | - | - | - |
| *Nuclearia sp.* | - | + | - | - | - |
| **Fungi** |  |  |  |  |  |
| *Allomyces macrogynus* | + | + | + | - | - |
| *Mortierella verticillata* | + | + | + | - | - |
| *Rozella allomycis* | + | + | + | - | - |
| *Spizellomyces punctatus* | + | + | + | - | - |

To better distinguish the RNase III-containing genes as either *Dicer* or *Drosha* we reconstructed a phylogeny with known Drosha and Dicer protein homologs from multiple animals and fungi (Figure 3). The resulting tree revealed a highly supported and distinct clades of Dicer (89% Maximum Likelihood (ML) bootstrap support and 1.0 Bayesian posterior probability (BP), hereafter ML/BP) and Drosha (81%/1.0). Within this framework, the ichthyosporean sequences were divided into several smaller clades that where phylogenetically related to both the Dicer and Drosha branches, supporting the gene annotation that both *Dicer* and *Drosha* genes are present in these species. In addition, there were several paralogs of both Dicer and Drosha among Ichthyosporeans, revealing several duplications of the genes, but unsupported phylogenetic relationships between these preclude an interpretation of the number of gene expansions and losses.

**Figure 3.**
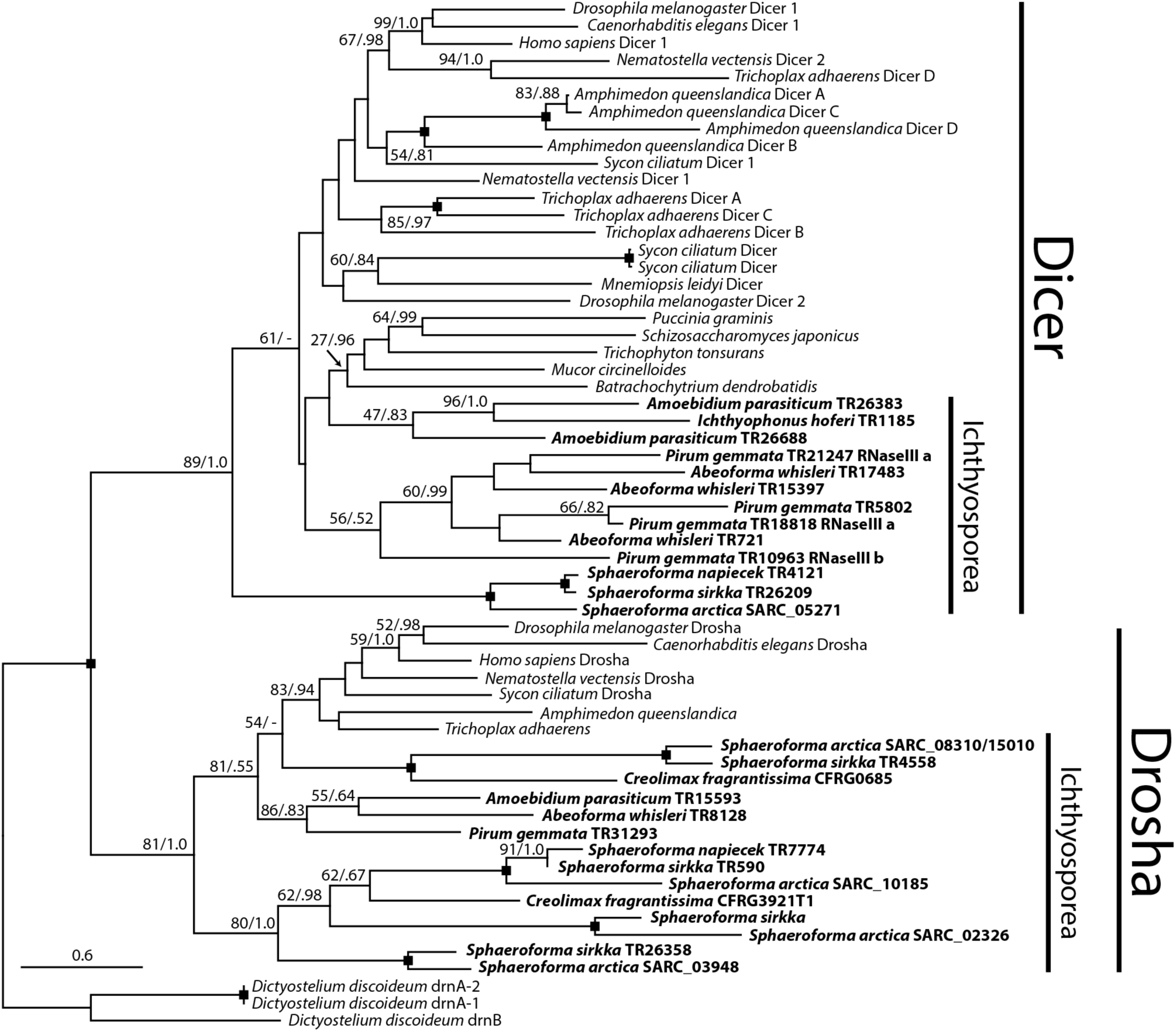
Phylogeny of Dicer and Drosha sequences across Opisthokonta. Phylogeny of Dicer and Drosha sequences of selected species covering Opisthokonta. The topology with the highest likelihood in a maximum likelihood (ML) framework is shown, with ML bootstrap and Bayesian posterior probabilities (BP) support values drawn onto the branching points (ML/BP). See Table S4 for accession numbers of gene sequences used in the analysis. Ichthyosporean species are drawn in bold font. Only support values above 50/.75 are shown, except at a few selected nodes. Maximum support (100/1.0) is indicated by a black square.

The phylogenetic analysis of the *Pasha* sequences identified with reciprocal Blast (Figure 4) confirmed that they are more closely related to the animal Pasha (86%/0.99 support for Pasha) than to HYL1, which was recently discovered also in basal animals (Moran *et al.*, 2013). The ichthyosporean sequences clustered as sister to animals in two separate clades, one consisting of *Sphaeroforma* and *Creolimax*(100%/1.0) and one of *Pirum gemmata*, *Abeoforma whisleri* and *Amoebidium parasiticum*(monophyly not supported), consistent with the mutual phylogenetic relationships within the Ichthyosporea.

**Fig 4.**
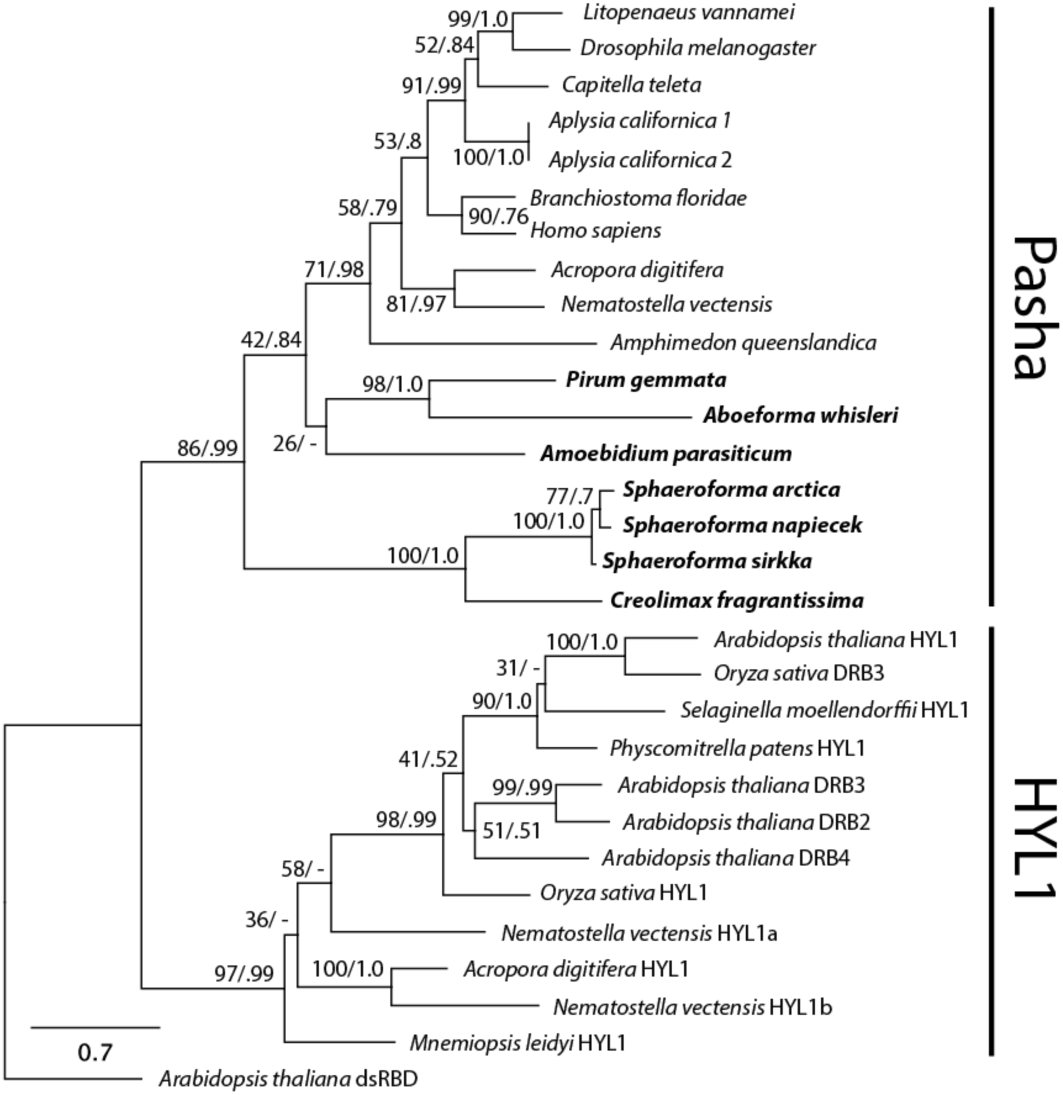
Pasha phylogeny. Phylogeny of Pasha and HYL1 sequences based on Moran *et al.*, 2013. Topology and support values created the same way as for Figure 2. See Table S5 for accession numbers of gene sequences used in the analysis. Ichthyosporean species are drawn in bold font.

## Discussion

### MicroRNAs are present in unicellular relatives of animals

Here we have sequenced the expressed small RNAs of three choanozoan species and for the first time shown evidence of miRNAs in unicellular relatives of animals. These miRNAs show all the characteristics of typical miRNAs, such as being transcribed from a structured miRNA precursor (pre-miRNA), processing by two consecutive Drosha and Dicer cuts that leave characteristic 2-nt 5’ and 3’ overhangs and 5’ homogeneous small RNA reads mapping to both the mature and star strands of the miRNAs. Four of the six novel miRNAs are highly conserved between the *Sphaeroforma* species we have investigated. Conservation has earlier been one of the criteria for defining miRNAs (Ambros *et al.*, 2003), and together with expression and the canonical folding structure this strongly indicate functional miRNA genes. Conservation of miRNAs across species has also been used as a criterion for the identification of novel miRNAs (Ambros *et al.*, 2003).

Only one of the newly detected miRNAs is within the typical range of animal miRNA loop sizes of up to 40 nts (Fromm *et al.*, 2015), while three others are only slightly above this observed threshold. However, two of the miRNAs have significantly larger loop sizes and resemble more those observed in animals with at least two separate Dicer genes, including plants, demosponges, pancrustacean arthropods, and flatworms (De Jong *et al.*, 2009; Gao *et al.*, 2014; Mukherjee *et al.*, 2013).

In an attempt to hone in on the function of these miRNAs we predicted their potential target genes in *S. arctica* which is the only species for which an annotation of the 3’-UTRs is available. Surprisingly, the predicted targeting mode of the miRNAs seems to be a mixture of animal- and plant-like mechanisms. For all the miRNAs we could easily identify several tens to more than hundred genes potentially targeted with an animal-type mechanism. And intriguingly, for the two miRNAs with longer loops (i.e. more plant-like) we could also identify several genes targeted in a plant-like manner. These targets were very different types of genes. Some of them have no recognizable homologs in public databases and could therefore be specific to *S. arctica*. One was homologous to an ABC transporter, which are ubiquitous among eukaryotes. And lastly, different paralogs of Zinc-finger transcription factors seem to be regulated by these miRNAs. These miRNAs may therefore have important regulatory functions by acting on transcription factors in ichthyosporeans.

Unfortunately, protocols and tools for assessing these regulatory functions by experimental approaches are obstructed by lack of appropriate molecular tools for *Sphaeroforma*(Suga and Ruiz-Trillo, 2013). Nevertheless, our results suggest that there is potentially a wide range of target genes in the genome of the unicellular Choanozoa reflecting the broad regulatory roles of the miRNA genes.

### Essential components of the miRNA protein machinery present in Ichthyosporea

The ichthyosporean miRNAs discovered here have all characteristics of being processed by double RNase III domain proteins because they have 2-nt 3’ overhangs and small RNAs that map to the mature and star strands of the pre-miRNAs. But until now *Dicer* or *Drosha*, as well as other key components of the animal miRNA biogenesis machinery, have not been identified in any ichthyosporean or other choanozoans (Maxwell *et al.*, 2012; Grimson *et al.*, 2008). We therefore re-examined available genome and transcriptome data from all choanozoans and identified *Pasha*, *XPO5* and *AGO* in all ichthyosperans investigated, as well as several genes that were either class II or III RNases which comprise *Dicer* and *Drosha* respectively (Nicholson, 2014; Kwon *et al.*, 2016). Strikingly, the phylogenetic annotation shows that all the ichthyosproreans harbor at last one copy, but more often several copies, of an RNase III gene which is more closely related to *Drosha* than *Dicer*. Drosha has previously only been found in animals and is believed to have evolved from Dicer, which is ubiquitous among eukaryotes (Maxwell *et al.*, 2012; Kwon *et al.*, 2016). In addition, homologs of the animal Dicer gene were identified in most ichthyosporeans.

The phylogenetic distribution of the *Dicer* and *Drosha* sequences reveal another pattern specific to the two closely related genera *Sphaeroforma* and *Creolimax*: Both possess two distinct subfamilies of *Drosha* that are exclusive to these groups. To our knowledge this is the first example of multiple copies of Drosha in a species.

### The Microprocessor complex evolved in ancient unicellular relatives of animals

Our findings demonstrate that miRNAs and a miRNA biogenesis pathway homologous to that in animals was innovated much earlier than in the last common ancestor of animals. This includes the animal-specific Microprocessor complex with Drosha and Pasha; in fact, these two genes are always found together. Our discovery of both Pasha and Drosha in all ichthyosporeans investigated suggests that the Microprocessor complex predates animals and was present already in the early history of opisthokonts. In contrast, no sign of the Microprocessor components was found in fungi or any basal Holomycota species. Altogether this suggest that the Microprocessor complex originated in Holozoa after the separation from Holomycota, but before Ichthyosporea diverged from the other choanozoan lineages.

This also implies that the absence of the Microprocessor complex, and miRNAs, in the deeply diverging comb jellies (ctenophores) must be a secondary loss. Hence it is not a primitive state argued to confirm their basal positon in the animal phylogeny (Maxwell *et al.*, 2012). Similarly, no member of the other two main Holozoa groups, the choanoflagellates and filastereans, show any signs of miRNAs or the Microprocessor machinery implying that they have independently lost these genes. Altogether therefore, the deep holozoan origin of the Microprocessor complex and miRNA genes has been followed by multiple independent losses.

Previously, nearly all classes of transcription factors, signaling and adhesion proteins in animals have been discovered among their unicellular relatives (see for instance Shalchian-Tabrizi *et al.*, 2008; Sebé-Pedrós and Ruiz-Trillo, 2016; Suga *et al.*, 2013; Richter and King, 2013; King *et al.*, 2008). Hence most of the genetic components required for complex multicellular organisms were innovated among unicellular sisters of animals. Here, we provide the first evidence that also the regulatory RNAs essential for animal gene regulation and development pre-date the origin of animals, suggesting that the last common ancestor of animals had miRNAs and the capacity of gene regulation by such small regulatory RNAs.

### Single origin and multiple losses of animal miRNAs

Animal miRNAs are believed to have originated at multiple occasions because sponge miRNAs are not conserved between silicisponges (demosponges + hexactinellids), calcisponges and homoscleromorphs, nor with other animals (Grimson *et al.*, 2008; Liew *et al.*, 2016; Robinson *et al.*, 2013; Wheeler *et al.*, 2009; Tarver *et al.*, 2012, 2015). According to this scenario, eumetazoan miRNAs are not homologs of sponge miRNAs and the last common ancestor of animals did not possess miRNA genes (Tarver *et al.*, 2015; Maxwell *et al.*, 2012; Moroz *et al.*, 2014).

However, the presence of the Microprocessor complex and the accompanying large variety of miRNA types in Ichthyosporea makes it likely that the ancestor of Holozoa and animals contained a variety of miRNA types. Hence, deviating animal miRNAs in the sponges may represent residuals of the ancient choanozoan genes instead of being genetic innovations. Accordingly, dissimilar miRNA types in the sponge lineages can be result of differential loss of the ancestral miRNAs.

The finding of conserved miRNAs outside of animals therefore changes how we view the evolution of miRNAs and their potential key role in the transition to complex multicellular animals. The striking similarities in the miRNA biogenesis machinery across the animal kingdom, which now also extends to include ichthyosporeans, argue for a common origin of the machinery and the capacity to process miRNAs that predates the animals. Hence the timing of the genetic innovations *per se* of the miRNAs in animals cannot longer be directly linked to the origin of multicellularity and complexity among animals.

## Materials and Methods

### Culturing and RNA sequencing

*S. arctica* JP610 cultures were grown on ATCC MAP medium at 16°C. The cells were lysed on a FastPrep system (MP Biomedicals, Santa Ana, CA, USA), followed by small RNA and total RNA isolation using the mirPremiere RNA kit (Sigma-Aldrich, St. Louis, MO, USA). For transcription start site (TSS) sequencing the total RNA was treated with Terminator exonuclease (Epicentre, Madison, WI, USA) and resistant mRNAs (i.e. carrying a 5'CAP) were sequenced (see supplementary information for more details). The RNA samples of *S. arctica* were sequenced on Illumina HiSeq2000 machine. Library preparation and sequencing was performed by vertis Biotechnologie AG (Freising, Germany).

*S. sirkka* B5 and *S. napiecek* B4 cultures were grown on Marine Broth (Difco BD, NJ, US) at 12°C. The cells were lysed and small RNAs isolated with the same method as for *S. arctica*. In addition, total RNA was isolated from both cultures for sequencing of the expressed mRNAs. miRNA libraries (20-40 nt fragment size) and mRNA libraries (300 nt paired-end) were prepared and sequenced on the Illumina MiSeq platform at the Norwegian Sequencing Centre.

### Mapping of RNA reads and miRNA detection

For *S. arctica*, mapping of all RNA reads was done against the 2012 version of the *S. arctica* genome, downloaded from the Broad Institute (http://www.broadinstitute.org). Also, 100 bp poly(A)-selected RNA Illumina reads from the SRX099331 and SRX099330 *S. arctica* experiments where downloaded from the National Center for Biotechnology Information (NCBI) Sequence Read Archive (SRA). All sequenced and downloaded RNA reads were trimmed for low quality nucleotides (phred score cutoff of 20) and sequencing adaptors using Trimmomatic v.0.30 (Lohse *et al.*, 2012), and trimmed for ‘N’ characters and poly(A)-tails using PrinSeq-lite v.0.20.3 Schmieder and Edwards, 2011. Additionally, only small RNAs reads between 18-24 nts were retained. For miRNA-detection, the miRDeep2* pipeline, an adapted version of mirDeep2 (Friedländer *et al.*, 2012) for the detection of longer hairpins as part of miRCandRef (https://hyperbrowser.uio.no/mircandref/; Fromm *et al.*, 2013) was used with default settings on the small RNA reads to first collapse identical reads, and subsequently map the collapsed reads to the *S. arctica* genome. TSS reads and poly(A)-selected reads were mapped to the *S. arctica* genome using TopHat v2.0.14 (Kim *et al.*, 2013) with default settings.

For *S. sirkka and S. napiecek*, small RNAs were trimmed using Trimmomatic v.0.36 to remove adapters and nucleotides with a quality < 28. Only reads longer than 19 nts were retained. The *S. sirkka* reads were mapped to the genome (downloaded from NCBI under accession LUCW01000000) using Bowtie v1.0.0 (Langmead et al. 2012) with the seed length set to 18 (-l 18) and maximum mismatches in the seed to 3 (-n 3). miRNAs were identified using mirDeep2* as for *S. arctica*. For *S. napiecek* there is no genome available so we could not run mirDeep2* for novel miRNA detection. Instead we mapped the expressed smallRNAs to the *de novo* assembled transcriptome (assembled using Trinity v2.0.6 (Grabherr *et al.*, 2011) with the – normalize_reads option set, otherwise default settings) with bowtie1 as described above. See Table S1 for sequencing and mapping statistics.

### miRNA target prediction

In order to identify targets for the *S. arctica* miRNAs, we extracted the 3’ UTR regions of all annotated *S. arctica* genes using the threeUTRsByTranscript() function in the GenomicFeatures R package (Lawrence *et al.*, 2013) and exporting them as a gff-file with the rtracklayer R package (Lawrence *et al.*, 2009). Fasta sequences of the 3’-UTRs were obtained using bedtools getfasta Quinlan and Hall, 2010. The TargetScan program (Lewis *et al.*, 2005) was used to identify potential miRNA targets in these sequences based on the mature strands of the miRNAs. As a first step, the targetscan_60.pl script was run to identify potential target sites. The resulting output was then submitted to the targetscan_60_context_scores.pl script in order to evaluate the miRNA-target interactions. Homology information was not included in the analysis. Only context+ scores < −0.3 were reported, similar to Jacobsen *et al.* (2013). Plant-like miRNA targets in the 3’-UTRs were identified using the TAPIR web server (Bonnet *et al.*, 2010). The program was used in the “precise” mode, with default parameters (mimic=0, score<=4, mfe_ratio>=0.7). Target genes were analyzed using Blastp and CD-search on the NCBI web servers.

To annotate the target genes identified by TAPIR, we blasted these against the NCBI nr and the Uniprot databases using Blastp searches on the NCBI web pages.

### Identification of genes related to the miRNA processing machinery

In order to search for the presence of genes involved in miRNA processing and function across Opisthokonta we searched available transcriptomes and genomes from a wide range of deep opisthokont species covering choanoflagellates, filastereans, ichthyosporeans and basal holomycotans (Table 2 and Table S3). For species which an assembled genome or transcriptome were not available, raw reads were downloaded from the NCBI SRA database, quality trimmed using Trimmomatic v0.35 (Bolger *et al.*, 2014) (minimum phred score 20-28 depending on read quality) and assembled using Spades v3.8.0 (Bankevich *et al.*, 2012) (with the –careful option, otherwise default settings) for genomic data and Trinity v2.0.6 (with the – normalize_reads option set, otherwise default settings) + Transdecoder v3.0.0 (Haas *et al.*, 2014) (TransDecoder.LongOrfs program with default settings) for transcriptomes where no reference genome was available and the TopHat v2.1.1 (Kim *et al.*, 2013) + Cufflinks v2.1.1 (Trapnell *et al.*, 2013) pipeline for transcriptomes when a reference genome was available.

As query genes we used Dicer, Drosha, Pasha, Argonaute (AGO) and Exportin 5 (XPO5) from *Homo sapiens*, *Drosophila melanogaster*, *Nematostella vectensis* and *Amphimedon queenslandica* and Dicer, AGO and XPO5 from the fungus *Neurospora crassa*. Accession numbers of the query genes are listed in Table S3.

Genes were identified using a reciprocal Blast strategy and domain identification. Blast was performed by searching the query sequences against each individual target genome/transcriptome using Blastp (Altschul *et al.*, 1990) (BLOSUM45 scoring matrix, min e-value 0.01 and max target hits 30). Each blast hit was then verified by reciprocal blast searches against a database consisting of the genomes and proteomes of the query organisms (i.e. *H. sapiens*, *D. melanogaster*, *N. vectensis*, *A. queenslandica*, *S. arctica* and *N. crassa*). All blast hits were sorted by increasing e-value. Only genes ranked as top hit in both reciprocal Blast runs were were retained. These hits were further verified by blasting against the UniProt database (same search parameters as above), and annotated as potential microRNA processing genes only when the UniProt search provided the same gene type match (as the query sequence) as the best hit. Further blast verification was usually performed against the GenBank nr database.

Domain annotation was performed using InterProScan and we defined miRNA-related genes on the basis of the identified domains as following: AGO: both PAZ and PIWI domains present, Dicer and Drosha: two RNase III domains present, Pasha: double stranded RNA-binding domain (dsRBD). XPO5 contains no conserved domains and was only identified with the reciprocal Blast strategy. For a few *Dicer* and *Drosha* sequences identified by reciprocal Blast, only a a single RNase III domain was identified with InterProScan. We therefore used an alignment strategy to verify the presence of a second domain (see Supplementary Material and Supplementary File 2). All blast searches and domain annotations were done using Geneious R9 (Kearse *et al.*, 2012), expect for the UniProt and GenBank blast searches which were performed on the UniProt and NCBI web sites.

### Phylogenetic annotation of miRNA processing proteins

A multiple sequence alignment containing known *Dicer* and *Drosha* sequences from animals, fungi and *Dictyostelium discoideum*, as well as the Dicer and Drosha sequences of ichthyosporeans identified in this study was generated using Mafft v7 Katoh and Toh, 2010. First, only full length Dicer and Drosha sequences were globally aligned using the G-INS-i algorithm and the BLOSUM45 scoring matrix, then shorter and incomplete sequences were added using the –add Fragments option. Finally, alignment columns containing > 90% gaps were removed. See Table S4 for list of accession numbers. Bayesian analysis was performed with PhyloBayes-MPI v1.5 (Lartillot *et al.*, 2009). Two chains were run with the parameters -gtr and -cat and stopped when the maxdiff was 0.18 and the meandiff 0.014 with a 15% burnin. Maximum likelihood (ML) analysis was run using RAxML v8.0.26 (Stamatakis, 2014) with the VT protein substitution model. The topology with the highest likelihood score out of 10 heuristic searches was selected as the final topology. Bootstrapping was carried out with 600 pseudo replicates under the same model. The values from the ML bootstrapping and the Bayesian posterior probabilities were added to the ML topology with the highest likelihood.

To investigate the evolutionary affiliation of the annotated Pasha sequences we created a multiple sequence alignment based on the study of Moran *et al.* (2013) consisting of Pasha and HYL1 (another double-stranded RNA binding protein which partner with Dicer) sequences (accession numbers are listed Table S5). The Pasha and HYL protein sequences were trimmed to only contain the double-stranded RNA binding domains (dsRBDs), with some remaining flanking positions, aligned together with the ichthyosporean Pasha candidates with Mafft (E-INS-i algorithm, BLOSUM45 scoring matrix on the mafft web-server: http://mafft.cbrc.jp/alignment/software/). The alignment was analyzed using ML and Bayesian analyses as described above. The ML analysis used the VT model and 500 pseudo-replicates and in the Bayesian the two chains had converged (burn-in 15%, maxdiff=0.04, meandiff=0.04). The values from the ML bootstrapping and the Bayesian posterior probabilities were added to the ML topology with the highest likelihood.

### Data accessibility

All smallRNA and transcriptome data generated in this study will be submitted to the publically available NCBI SRA database upon acceptance of the manuscript. The novel miRNA candidates will be submitted to MirGeneDB.org and sequences of the miRNA processing genes in will be submitted to NCBI GenBank. In addition, sequence alignments used in the phylogenetic analyses will be made available on the Dryad database (datadryad.org) and the Bioportal (www.bioportal.no).

## Acknowledgments

We are grateful to Iñaki Ruiz-Trillo for providing the *S. arctica* culture and to Brandon Hassett for providing the *S. sirkka* and *S. napiecek* cultures. We thank Notur for providing computational resources.

The postdoc grants (Nr. 213703 and 240284) to JB was funded by the Norwegian Research Council, while BF is supported by South-Eastern Norway Regional Health Authority grant 2014041. PhD fellowship for RSN and research grant to KS-T was funded by the Molecular Life Science board at University of Oslo.

## Author contributions

JB participated in the study design, took part in all the data analyses, made figures and drafted the manuscript. RSN participated in the study design, cultured and isolated RNA from *S. arctica*, analyzed the *S. arctica* smallRNAs and the miRNA pathway genes and wrote the initial manuscript draft. AABH cultured and isolated RNA from *S. sirkka* and *S. napiecek*, assembled novel transcriptomes and genomes, analysed the smallRNAs, developed the reciprocal Blast pipeline and commented on the manuscript. BF analyzed the smallRNA data, identified and annotated miRNAs, provided critical evaluation of the miRNA structures, participated in figure design and commented on the manuscript. PEG participated in the study design, provided critical discussion on miRNA function and commented on the manuscript. KST participated in the study design, evaluated all the data analyses and figures and contributed on the initial and final manuscripts. All authors have read and approved the final manuscript.

## Supplementary information

### Transcription start site (TSS) sequencing of Sphaeroforma arctica

*S. arctica* cells were grown and lysed as described above; RNA was isolated using the total RNA protocol of the mirPremiere RNA kit. The RNA samples were fragmented with ultrasound and then treated with T4 polynucleotide kinase (PNK). The RNA species which carry a 5' mono-phosphate were degraded using Terminator exonuclease (Epicentre, Madison, WI, USA). The exonuclease resistant mRNA (primary transcripts with 5'CAP) were poly(A)-tailed using poly(A) polymerase. The sample was then split into two parts; one half (5' +TAP sample) was treated with tobacco acid pyrophosphatase (Epicentre) to degrade 5'CAP ends to 5'P, while the other half (5' -TAP sample) was left untreated. Afterwards, an RNA adapter was ligated to the 5'-phosphate of the RNA fragments. First-strand cDNA synthesis was performed using an oligo(dT)-adapter primer and M-MLV reverse transcriptase. The resulting cDNA was PCR-amplified to about 20-30 ng/μl using a high fidelity DNA polymerase (13 and 14 cycles for the treated and untreated samples, respectively). The cDNA was purified using the Agencourt AMPure XP kit (Beckman Coulter Genomics). Sequencing was performed on an Illumina HiSeq2000 machine. Library preparation and sequencing was performed by Vertis Biotechnologie AG.

### Identification of Dicer and Drosha RNase III domains in Ichthyosporeans

Only two gene families contain double RNase III domains and these comprise the Drosha and Dicer and genes (i.e. class II and III of the RNAase III gene family; Kwon et al). For some putative ichthyosporean *Dicer*/*Drosha* homologs, conventional domain search tools such as Pfam or CD-search identified only one RNase III domain located in the C-terminal region. We therefore aligned the sequences to the RNase III a and b domains of other animal and fungal Dicers and Drosha proteins, as well as the bacterial *Aquifex aeolicus* RNase III domain (names and accession numbers in Supplementary File 2). The alignment was done by splitting the sequences into parts consisting of only the RNaseIII a or b domain. For sequences without an annotated RNaseIII domain these putative domains were identified by aligning the sequence to the annotated domains of the *H. sapiens* and *N. vectensis* Dicer and Drosha sequences. Then all RNaseIII a and b domains were aligned together. All alignments were done using Mafft v.7 Katoh and Toh, 2010 with the L-INS-I algorithm with the BLOSUM45 scoring matrix. The alignment revealed that the RNase III a domain is much more divergent than the RNase III b domain, confirming earlier reports (Shi *et al.*, 2006). Nevertheless it shows clear homology of the RNase III b domain for all ichthyosporean Dicer/Drosha sequences, and reveal homologous regions in the RNase III a domain, including residues in the highly important signature motif (Ji, 2008).

**Table S1.** Number of RNA-Seq reads before and after quality trimming and mapping.

|  | Raw reads | After trimming | Mapped reads |
| --- | --- | --- | --- |
| *Sphaeroforma arctica* |  |  |  |
| small RNA | 10,128,170 | 2,309,997 | 2,125,525 |
| TSS (tap +) | 26,383,940 | 25,792,655 | 12,759,092 |
| TSS (tap -) | 17,898,227 | 17,527,249 | 3,754,208 |
| mRNA | 28,487,927 | 23,961,010 | 21,361,551 |
| *Sphaeroforma sirkka* |  |  |  |
| small RNA | 6,464,007 | 4,379,752 | 2,787,653 |
| mRNA | 14,258,902 | 13,854,020 | 11,242,128 |
| *Sphaeroforma napiecek* |  |  |  |
| small RNA | 6,569,325 | 4,696,713 | 4,113,599* |
| mRNA | 14,239,644 | 13,865,534 | NA |

\* Mapped to the assembled transcriptome

**Figure S1.**
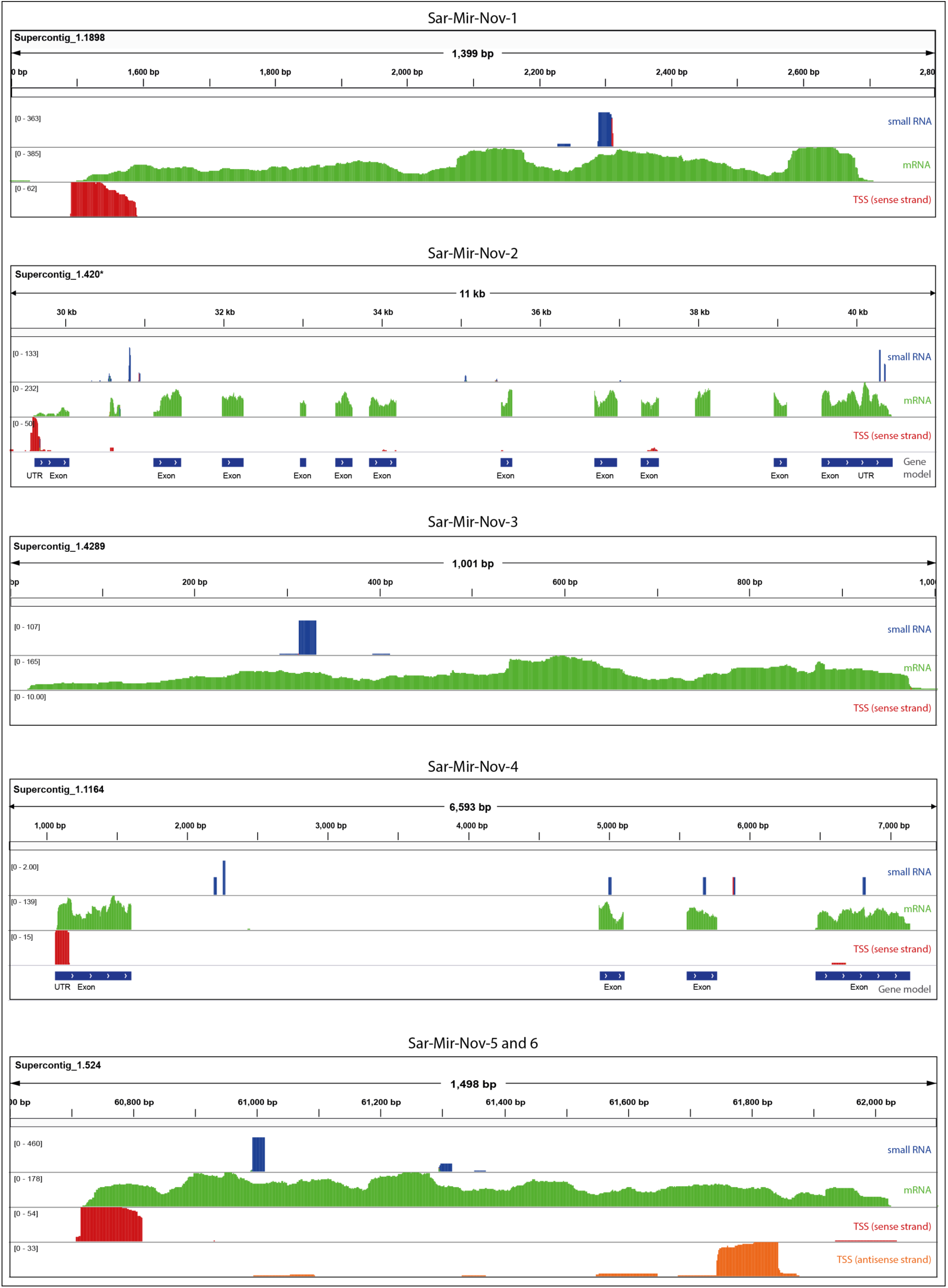
Genomic localization and expression of *S. arctica* miRNAs. Mapping of smallRNAs (blue track), mRNAs (green track) and transcription start site (TSS) reads (red and orange tracks) at the genomic regions surrounding the miRNAs in *Sphaeroforma arctica*. *For Supercontig_1.420 we merged Supercontig_1.3651 which contain the middle domain of AGO and are expressed together in the transcriptome. Note that Sar-Mir-Nov-3 is lacking a TSS but this is probably an artifact of the genome assembly (the contig is very short and the mRNAs map almost from beginning to the end).

**Table S2.** Plant-like targeting in *S. arctica*. Genes in *Sphaeroforma arctica* targeted by the novel miRNAs in a plant-like manner. Genes annotated using Geneious R9 by running InterProScan, and Blastp searches against NCBI RefSeq database (with BLOSUM45 scoring matrix) and the UniprotKB/Swiss-prot database. Only Blast hits with an e-value < 0.05 are shown. And in cases where the top his was an un-annotated gene, the best hitting annotate gene is shown. Hits against *S. arctica* genes in RefSeq are not reported.

| miRNA | *S. arctica* target gene | InterProScan | RefSeq (e-value) | Uniprot/Swissprot (e-value) |
| --- | --- | --- | --- | --- |
| Sar-Mir-Nov-1 | No targets |  |  |  |
| Sar-Mir-Nov-2 | No targets |  |  |  |
| Sar-Mir-Nov-3 | SARC_11150 | ABC transporter | ATP binding cassette transporter (1.38e-146) | ABC transporter G family member 22 (1.39e-79) |
| Sar-Mir-Nov-4 | SARC_09903 | Not annotated | No hits | RED1 (4.84e-02) |
| Sar-Mir-Nov-5 | SARC_16580 | Zinc finger containing protein | No hits | Krueppel like factor 16 (1.08e-21) |
|  | SARC_15089 | Zinc finger containing protein | No hits | Zinc finger protein 76 (1.45e-51) |
|  | SARC_03864 | Zinc finger containing protein | No hits | Zinc finger protein (1.03e-21) |
| Sar-Mir-Nov-6 | SARC_16580 | Zinc finger containing protein | No hits | Krueppel like factor 16 (1.08e-21) |
|  | SARC_15089 | Zinc finger containing protein | No hits | Zinc finger protein 76 (1.45e-51) |
|  | SARC_03864 | Zinc finger containing protein | No hits | Zinc finger protein (1.03e-21) |

**Table S3.** Genes used as queries for the detection of miRNA processing genes

| Species | Gene name | Accession number |
| --- | --- | --- |
| *Homo sapiens* | Argonaute-1 | NP_001304051 |
|  | Argonaute-2 | NP_036286 |
|  | Argonaute-3 | NP_079128 |
|  | Argonaute-4 | NP_060099 |
|  | Exportin 5 | NP_065801 |
|  | Dicer 1 | NP_085124 |
|  | Ribonuclease 3 | NP_037367 |
|  | DGCR8 | NP_073557 |
|  | DGCR8 | NP_001177255 |
| *Drosophila melanogaster* | Argonaute-1 | NP_725341 |
|  | Argonaute-2 | ABB54719 |
|  | Argonaute-3 | ABO27430 |
|  | RanBP21 | AF222746 |
|  | Dicer 1 | NP_524453 |
|  | Dicer 2 | NP_523778 |
|  | Drosha | NP_477436 |
|  | Pasha | NP_651879 |
|  | Pasha | NP_001263149 |
| *Nematostella vectensis* | Argonaute-1 | AGW15594 |
|  | Argonaute-2 | AGW15595 |
|  | NEMVEDRAFT_v1g144913 | XP_001621438 |
|  | Dicer 1 | AGW15597 |
|  | Dicer 2 | AGW15596 |
|  | Drosha | AGW15598 |
|  | Pasha/DGCR8 | AGW15599 |
| *Amphimedon queenslandica* | piwi-like protein 1 | XP_011409849 |
|  | argonaute-2-like | XP_003385988 |
|  | exportin-5-like | XP_011402610 |
|  | Dicer-like | XP_003384628 |
|  | Dicer-like | XP_011402849 |
|  | Dicer-like | XP_003391904 |
|  | Dicer-like | XP_003384628 |
|  | Ribonuclease-3-like | XP_011404936 |
| *Neurospora crassa* | sms-2 | XP_958586 |
|  | QDE2 | AAF43641 |
|  | NCU02387 | XP_959707 |
|  | Dicer-like protein 1 | XP_961898 |
|  | Dicer-like-2 | XP_963538 |

**Table S4.** Genes used in the Dicer and Drosha phylogeny

| Species | Accession number | Comment |
| --- | --- | --- |
| *Amoebidium parasiticum* | TR26383 | *de novo* assembly in this study |
| *Amoebidium parasiticum* | TR26688 | *de novo* assembly in this study |
| *Amoebidium parasiticum* | TR15593 | *de novo* assembly in this study |
| *Amphimedon queenslandica* | XP_003384628 |  |
| *Amphimedon queenslandica* | XP_003385258 |  |
| *Amphimedon queenslandica* | XP_003391904 |  |
| *Amphimedon queenslandica* | contig13288 | From Mukherjee et al. 2013 |
| *Amphimedon queenslandica* | XP_011404936 |  |
| *Aboeforma whisleri* | TR721 | *de novo* assembly in this study |
| *Aboeforma whisleri* | TR15397 | *de novo* assembly in this study |
| *Aboeforma whisleri* | TR17483 | *de novo* assembly in this study |
| *Aboeforma whisleri* | TR8128 | *de novo* assembly in this study |
| *Batrachochytrium dendrobatidis* | EGF82843 |  |
| *Caenorhabditis elegans* | NP_498761 |  |
| *Caenorhabditis elegans* | NP_492599 |  |
| *Creolimax fragrantissima* | CFRG0685 |  |
| *Creolimax fragrantissima* | CFRG3921 |  |
| *Drosophila melanogaster* | NP_524453 |  |
| *Drosophila melanogaster* | NP_523778 |  |
| *Drosophila melanogaster* | AAD31170 |  |
| *Dictyostelium discoideum* | EAL70472 |  |
| *Dictyostelium discoideum* | EAL71163 |  |
| *Dictyostelium discoideum* | EAL73658 |  |
| *Homo sapiens* | NP_085124 |  |
| *Homo sapiens* | Q9NRR4 |  |
| *Ichthyophonus hoferi* | TR1185 | *de novo* assembly in this study |
| *Mucor circinelloides* | CAK32533 |  |
| *Mnemiopsis leidyi* | AFX00003 |  |
| *Nematostella vectensis* | ABZ10549 |  |
| *Nematostella vectensis* | ABZ10551 |  |
| *Nematostella vectensis* | AGW15598 |  |
| *Puccinia graminis* | EFP86699 |  |
| *Pirum gemmata* | TR5802 | *de novo* assembly in this study |
| *Pirum gemmata* | TR10963 | *de novo* assembly in this study |
| *Pirum gemmata* | TR18818 | *de novo* assembly in this study |
| *Pirum gemmata* | TR21247 | *de novo* assembly in this study |
| *Pirum gemmata* | TR31293 | *de novo* assembly in this study |
| *Sphaeroforma arctica* | SARC_08310 and SARC_15010 | Genes connected by mRNA reads |
| *Sphaeroforma arctica* | SARC_02326 |  |
| *Sphaeroforma arctica* | SARC_03948 |  |
| *Sphaeroforma arctica* | SARC_05271 |  |
| *Sphaeroforma arctica* | SARC_10185 |  |
| *Schizosaccharomyces japonicus* | XP_002174825 |  |
| *Sphaeroforma napiecek* | TR4121 | *de novo* assembly in this study |
| *Sphaeroforma napiecek* | TR7774 | *de novo* assembly in this study |
| *Sphaeroforma sirkka* | TR590 | *de novo* assembly in this study |
| *Sphaeroforma sirkka* | TR4558 | *de novo* assembly in this study |
| *Sphaeroforma sirkka* | TR26209 | *de novo* assembly in this study |
| *Sphaeroforma sirkka* | TR26358 | *de novo* assembly in this study |
| *Sphaeroforma sirkka* | NA | From LUCW01036240 and<br>LUCW01041741 genome contigs |
| *Sycon ciliatum* | scpid37237 | From Compagen.org |
| *Sycon ciliatum* | scpid10519 | From Compagen.org |
| *Sycon ciliatum* | scpid10786 | From Compagen.org |
| *Sycon ciliatum* | scpid5641 | From Compagen.org |
| *Trichoplax adhaerens* | XP_002108754 |  |
| *Trichoplax adhaerens* | XP_002107798 |  |
| *Trichoplax adhaerens* | XP_002107959 |  |
| *Trichoplax adhaerens* | XP_002108012 |  |
| *Trichoplax adhaerens* | XP_002109903 |  |
| *Trichophyton tonsurans* | EGD96414 |  |

**Table S5.** Genes used in the Pasha phylogeny

| Species | Accession number | Note |
| --- | --- | --- |
| *Aplysia californica* | XP_005109075 |  |
| *Aplysia californica* | XP_005109080 |  |
| *Acropora digitifera* | XP_015764940 |  |
| *Acropora digitifera* | XP_015769232 |  |
| *Amoebidium parasiticum* | TR13520 | *de novo* assembly in this study |
| *Amphimedon queenslandica* | XP_011405426 |  |
| *Arabidopsis thaliana* | Q9SP32 |  |
| *Arabidopsis thaliana* | Q9SKN2 |  |
| *Arabidopsis thaliana* | Q9LJF5 |  |
| *Arabidopsis thaliana* | Q8H1D4 |  |
| *Arabidopsis thaliana* | O04492 |  |
| *Aboeforma whisleri* | TR9624 | *de novo* assembly in this study |
| *Amoebidium parasiticum* | TR13520 |  |
| *Branchiostoma floridae* | XP_002589316 |  |
| *Creolimax fragrantissima* | CFRG6281T1 |  |
| *Capitella teleta* | ELT95530 |  |
| *Drosophila melanogaster* | NP_651879 |  |
| *Homo sapiens* | NP_073557 |  |
| *Litopenaeus vannamei* | AEI83216 |  |
| *Mnemiopsis leidyi* | ML025024a | Available from Mnemiopsis genome project portal |
| *Nematostella vectensis* | AGW15600 |  |
| *Nematostella vectensis* | AGW15601 |  |
| *Nematostella vectensis* | XP_001636661 |  |
| *Oryza sativa* | Q0DJA3 |  |
| *Oryza sativa* | Q5N8Z0 |  |
| *Pirum gemmata* | TR7874 | *de novo* assembly in this study |
| *Physcomitrella patens* | XP_001782891 |  |
| *Sphaeroforma arctica* | SARC_09010 |  |
| *Selaginella moellendorffii* | XP_002977089 |  |
| *Sphaeroforma napiecek* | TR7380 | *de novo* assembly in this study |
| *Sphaeroforma sirkka* | TR3181 | *de novo* assembly in this study |

## References

Agarwal V, Bell GW, Nam JW, Bartel DP. (2015). Predicting effective microRNA target sites in mammalian mRNAs. Elife 4: 1–38.

Altschul SF, Gish W, Miller W, Myers EW, Lipman DJ. (1990). Basic Local Alignment Search Tool. J Mol Biol 215: 403–410.

Ambros V, Bartel B, Bartel DP. (2003). A uniform system for micro RNA annotation. Rna 9: 277–279.

Axtell MJ, Westholm JO, Lai EC. (2011). Vive la différence: biogenesis and evolution of microRNAs in plants and animals. Genome Biol 12: 221.

Bankevich A, Nurk S, Antipov D, Gurevich AA, Dvorkin M, Kulikov AS, et al. (2012). SPAdes: A New Genome Assembly Algorithm and Its Applications to Single-Cell Sequencing. J Comput Biol 19: 455–477.

Bartel DP. (2009). MicroRNAs: Target Recognition and Regulatory Functions. Cell 136: 215–233.

Berezikov E. (2011). Evolution of microRNA diversity and regulation in animals. Nat Rev Genet 12: 846–860.

Bolger AM, Lohse M, Usadel B. (2014). Trimmomatic: A flexible trimmer for Illumina sequence data. Bioinformatics 30: 2114–2120.

Bonnet E, He Y, Billiau K, van de Peer Y. (2010). TAPIR, a web server for the prediction of plant microRNA targets, including target mimics. Bioinformatics 26: 1566–1568.

Cerutti H, Casas-Mollano JA. (2006). On the origin and functions of RNA-mediated silencing: from protists to man. Curr Genet 50: 81–99.

Erwin DH, Laflamme M, Tweedt SM, Sperling EA, Pisani D, Peterson KJ. (2011). The Cambrian Conundrum: Early Divergence and Later Ecological Success in the Early History of Animals. Science (80-) 334: 1091–1097.

Friedländer MR, Mackowiak SD, Li N, Chen W, Rajewsky N. (2012). miRDeep2 accurately identifies known and hundreds of novel microRNA genes in seven animal clades. Nucleic Acids Res 40: 37–52.

Fromm B. (2016). microRNA Discovery and Expression Analysis in Animals. In: Field Guidelines for Genetic Experimental Designs in High-Throughput Sequencing, Aransay, AM & Trueba, L (eds), Springer International Publishing, pp 121–142.

Fromm B, Billipp T, Peck LE, Johansen M, Tarver JE, King BL, et al. (2015). A Uniform System for the Annotation of Vertebrate Microrna Genes and the Evolution of the Human Micrornaome. Annu Rev Genet 1017–1026.

Fromm B, Worren MM, Hahn C, Hovig E, Bachmann L. (2013). Substantial Loss of Conserved and Gain of Novel MicroRNA Families in Flatworms. Mol Biol Evol 30: 2619–2628.

Gao Z, Wang M, Blair D, Zheng Y, Dou Y. (2014). Phylogenetic analysis of the endoribonuclease Dicer family. PLoS One 9: 1–7.

Grabherr MG, Haas BJ, Yassour M, Levin JZ, Thompson DA, Amit I, et al. (2011). Full-length transcriptome assembly from RNA-Seq data without a reference genome. Nat Biotechnol 29: 644–652.

Grimson A, Srivastava M, Fahey B, Woodcroft BJ, Chiang HR, King N, et al. (2008). Early origins and evolution of microRNAs and Piwi-interacting RNAs in animals. Nature 455: 1193–7.

Haas BJ, Papanicolaou A, Yassour M, Grabherr M, Philip D, Bowden J, et al. (2014). De novo transcript sequence reconstruction from RNA-Seq: reference generation and analysis with Trinity. Nat Protoc 8: 1–43.

Han J, Lee Y, Yeom KH, Nam JW, Heo I, Rhee JK, et al. (2006). Molecular Basis for the Recognition of Primary microRNAs by the Drosha-DGCR8 Complex. Cell 125: 887–901.

Jacobsen A, Silber J, Harinath G, Huse JT, Schultz N, Sander C. (2013). Analysis of microRNA-target interactions across diverse cancer types. Nat Struct Mol Biol 20: 1325–32.

De Jong D, Eitel M, Jakob W, Osigus H-JJ, Hadrys H, Desalle R, et al. (2009). Multiple dicer genes in the early-diverging metazoa. Mol Biol Evol 26: 1333–40.

Katoh K, Toh H. (2010). Parallelization of the MAFFT multiple sequence alignment program. Bioinformatics 26: 1899–900.

Kearse M, Moir R, Wilson A, Stones-Havas S, Cheung M, Sturrock S, et al. (2012). Geneious Basic: An integrated and extendable desktop software platform for the organization and analysis of sequence data. Bioinformatics 28: 1647–1649.

Kim D, Pertea G, Trapnell C, Pimentel H, Kelley R, Salzberg SL. (2013). TopHat2: accurate alignment of transcriptomes in the presence of insertions, deletions and gene fusions. Genome Biol 14: R36.

King N, Westbrook MJ, Young SL, Kuo A, Abedin M, Chapman J, et al. (2008). The genome of the choanoflagellate Monosiga brevicollis and the origin of metazoans. Nature 451: 783–8.

Kosik KS. (2010). MicroRNAs and cellular phenotypy. Cell 143: 21–26.

Kwon SC, Nguyen TA, Choi Y-G, Jo MH, Hohng S, Kim VN, et al. (2016). Structure of Human DROSHA. Cell 164: 81–90.

Lartillot N, Lepage T, Blanquart S. (2009). PhyloBayes 3: A Bayesian software package for phylogenetic reconstruction and molecular dating. Bioinformatics 25: 2286–2288.

Lawrence M, Gentleman R, Carey V. (2009). rtracklayer: An R package for interfacing with genome browsers. Bioinformatics 25: 1841–1842.

Lawrence M, Huber W, Pagès H, Aboyoun P, Carlson M, Gentleman R, et al. (2013). Software for Computing and Annotating Genomic Ranges. PLoS Comput Biol 9: 1–10.

Lewis BP, Burge CB, Bartel DP. (2005). Conserved seed pairing, often flanked by adenosines, indicates that thousands of human genes are microRNA targets. Cell 120: 15–20.

Liew YJ, Ryu T, Aranda M, Ravasi T. (2016). miRNA Repertoires of Demosponges Stylissa carteri and Xestospongia testudinaria. PLoS One 11: e0149080.

Lohse M, Bolger AM, Nagel A, Fernie AR, Lunn JE, Stitt M, et al. (2012). RobiNA: A user-friendly, integrated software solution for RNA-Seq-based transcriptomics. Nucleic Acids Res 40: 622–627.

Mattick JSJ. (2004). RNA regulation: a new genetics? Nat Rev Genet 5: 1662–1666.

Maxwell EK, Ryan JF, Schnitzler CE, Browne WE, Baxevanis AD. (2012). MicroRNAs and essential components of the microRNA processing machinery are not encoded in the genome of the ctenophore Mnemiopsis leidyi. BMC Genomics 13. doi:10.1186/1471-2164-13-714.

Mello CC, Conte D. (2004). Revealing the world of RNA interference. Nature 431: 338–342.

Moran Y, Praher D, Fredman D, Technau U. (2013). The evolution of MicroRNA pathway protein components in Cnidaria. Mol Biol Evol 30: 2541–2552.

Moroz LL, Kocot KM, Citarella MR, Dosung S, Norekian TP, Povolotskaya IS, et al. (2014). The ctenophore genome and the evolutionary origins of neural systems. Nature. doi:10.1038/nature13400.

Mukherjee K, Campos H, Kolaczkowski B. (2013). Evolution of animal and plant dicers: early parallel duplications and recurrent adaptation of antiviral RNA binding in plants. Mol Biol Evol 30: 627–41.

Nicholson AW. (2014). Ribonuclease III mechanisms of double-stranded RNA cleavage. Wiley Interdiscip Rev RNA 5: 31–48.

Philippe H, Brinkmann H, Copley RR, Moroz LL, Nakano H, Poustka AJ, et al. (2011). Acoelomorph flatworms are deuterostomes related to Xenoturbella. Nature 470: 255–258.

Quinlan AR, Hall IM. (2010). BEDTools: A flexible suite of utilities for comparing genomic features. Bioinformatics 26: 841–842.

Richter DJ, King N. (2013). The Genomic and Cellular Foundations of Animal Origins. Annu Rev Genet. doi:10.1146/annurev-genet-111212-133456.

Robinson JM, Sperling EA, Bergum B, Adamski M, Nichols SA, Adamska M, et al. (2013). The Identification of MicroRNAs in Calcisponges: Independent Evolution of MicroRNAs in Basal Metazoans. J Exp Zool Part B Mol Dev Evol 320: 84–93.

Schmieder R, Edwards R. (2011). Quality control and preprocessing of metagenomic datasets. Bioinformatics 27: 863–864.

Sebé-Pedrós A, Ruiz-Trillo I. (2016). Evolution and Classification of the T-Box Transcription Factor Family. Curr Top Dev Biol 1–26.

Shabalina SA, Koonin E V. (2016). Origins and evolution of eukaryotic RNA interference. Trends Ecol Evol 23: 578–587.

Shalchian-Tabrizi K, Minge M A, Espelund M, Orr R, Ruden T, Jakobsen KS, et al. (2008). Multigene phylogeny of choanozoa and the origin of animals. PLoS One 3: e2098.

Stamatakis A. (2014). RAxML version 8: A tool for phylogenetic analysis and post-analysis of large phylogenies. Bioinformatics 30: 1312–1313.

Suga H, Chen Z, de Mendoza A, Sebé-Pedrós A, Brown MW, Kramer E, et al. (2013). The Capsaspora genome reveals a complex unicellular prehistory of animals. Nat Commun 4: 2325.

Suga H, Ruiz-Trillo I. (2013). Development of ichthyosporeans sheds light on the origin of metazoan multicellularity. Dev Biol 277: 284–292.

Tarver JE, Cormier A, Pinzón N, Taylor RS, Carré W, Strittmatter M, et al. (2015). microRNAs and the evolution of complex multicellularity: Identification of a large, diverse complement of microRNAs in the brown alga Ectocarpus. Nucleic Acids Res 43: 6384–6398.

Tarver JE, Donoghue PCJ, Peterson KJ. (2012). Do miRNAs have a deep evolutionary history? BioEssays 34: 857–66.

Thompson EA, Roeder GS. (1989). Expression and DNA sequence of RED1, a gene required for meiosis I chromosome segregation in yeast. Mol Gen Genet 218: 293–301.

Trapnell C, Hendrickson DG, Sauvageau M, Goff L, Rinn JL, Pachter L. (2013). Differential analysis of gene regulation at transcript resolution with RNA-seq. Nat Biotechnol 31: 46–53.

Wheeler BM, Heimberg AM, Moy VN, Sperling EA, Holstein TW, Heber S, et al. (2009). The deep evolution of metazoan microRNAs. Evol Dev 11: 50–68.

## References

Ji X. (2008). The mechanism of RNase III action: How Dicer dices. Curr Top Microbiol Immunol 320: 99–116.

Shi H, Tschudi C, Ullu E. (2006). An unusual Dicer-like1 protein fuels the RNA interference pathway in Trypanosoma brucei. RNA 12: 2063–2072.

